# Characterization of the pulmonary immune response induced by a highly protective tuberculosis vaccine using latent and acute infection mouse models

**DOI:** 10.64898/2025.12.04.692326

**Authors:** Steven C. Derrick, Amy Yang, Siobhan Cowley

## Abstract

We utilized a mouse tuberculosis (TB) latency model to evaluate multiple candidate TB vaccines for their ability to prevent reactivation of a latent infection. Among the most promising vaccine regimens tested was BCG formulated in adjuvant (Adj) (dimethyl dioctadecyl-ammonium bromide (DDA) plus D-(+)-Trehalose 6,6’-Dibehenate (TDB)) with rEsat-6 delivered subcutaneously (SC) followed by an intranasal (IN) administration of an adenovirus construct expressing a fusion of Esat-6 (E6) and antigen-85B (Ag85B) (AdE6-85B) before an aerosol *M. tuberculosis* challenge. We designated this vaccine regimen as BAA. BAA consistently prevented reactivation of 75 - 100% of immunized animals and was also highly and significantly protective against an acute aerosol infection with a consistent 2 – 3 log_10_ mycobacterial CFU reduction in the lungs relative to nonimmunized mice (Naïve). Likewise, the BAA vaccine was significantly more protective than BCG or BCG+Adj controls. Interestingly, we found that pre-challenge frequencies of CD4^+^ tissue resident memory (T_RM_) T cells (CD69^+^PD-1^+^CXCR3^+^), and CD4^+^ populations bearing CD153 and P2X7R, which are markers for protection, were significantly elevated in the lungs of mice immunized with the BAA vaccine relative to control groups. Additionally, we found significantly higher frequencies of multifunctional CD4^+^ T cells from infected lungs expressing both IL-17A and TGFβ or IL-17A, TGFβ and IFN-γ than in control groups. These findings suggest that vaccine regimens that establish a population of CD4^+^ T_RM_ cells in the lungs prior to infection and populations of multifunctional CD4^+^ T cells after infection may help control an acute pulmonary infection and prevent progression to active disease.

**IMPORTANCE:** To help curtail the TB epidemic, a new vaccine should prevent progression from a latent infection to active disease. Correlates of protective immunity, however, are presently unclear, which impedes the development of an improved TB vaccine. Hence, we tested different vaccines for their ability to prevent reactivation using a mouse latency model and identified a highly efficacious formulation using this model and, also, after testing using an acute aerosol infection model. We then examined the pulmonary immune responses induced by this vaccine both before and after an aerosol challenge and identified immune markers as well as populations of multifunctional and tissue resident memory T cells that may serve as correlates of vaccine efficacy against progression to active TB disease and control of an acute infection.

## INTRODUCTION

Tuberculosis (TB) remains an immense public health concern with around one-fourth of the world’s population is reported to be infected with *Mycobacterium tuberculosis* (1). The World Health Organization estimated a total of 1.25 million deaths including 161,000 among HIV-infected population (1.6 million total) were attributed to TB with an estimated 10.8 million people acquired TB disease in 2023 (1). Importantly, control of this epidemic has been confounded by the emergence of multiple-drug-resistant (MDR) and extensively drug-resistant (XDR) *M. tuberculosis* strains which often limit treatment options and make appropriate medical interventions challenging. Globally, there were approximately 176,000 MDR and 29,000 XDR TB cases in 2023 (1).

The only licensed vaccine against TB, *M. bovis* BCG, is administered to approximately 100 million children annually and is effective in reducing cases of severe disseminated tuberculosis (TB meningitis and miliary TB) in children (2, 3). However, the effectiveness of BCG vaccine in preventing the most contagious and prevalent form of disease, pulmonary TB, is unsatisfactory. BCG-induced protection against TB has been highly variable with protective efficacies ranging from 0 - 80% in numerous clinical trials (2, 4). Furthermore, protection induced by BCG vaccination is often not highly persistent and a substantial waning of the protective responses is generally seen during the first decade after immunization (5). Given the severity of the global TB problem and the sub-optimal effectiveness of BCG immunization, an improved TB vaccination strategy is urgently needed. However, a major impediment to the development of a more effective vaccine is that correlates of protective immunity to tuberculosis remain ill-defined.

We and others previously found that immune responses elicited by vaccinations with BCG formulated in DDA/TDB adjuvant (BCG+Adj) provided significantly better control of *M. tuberculosis* growth in the lungs of mice after aerosol challenge than nonadjuvanted BCG, and that immunization with the adjuvant alone was not protective (6–9). We subsequently showed that IL-17A expression from CD4^+^ T cells was necessary for the enhanced protection associated with the BCG+Adj vaccine (10). For the work described here, we used a TB latency mouse model to screen candidate vaccines capable of reducing reactivation rates so that we could subsequently evaluate the pulmonary immune response in the immunized animals with the aim of gaining a better understanding of what constitutes a protective immune response against progression to active disease. Since the BCG+Adj vaccine previously demonstrated superior protection using an acute infection mouse model, we focused our efforts on different vaccine regimens incorporating BCG formulated in DDA/TDB adjuvant. We also focused on the induction of different immune markers known to be associated with protection against an acute infection as well as a vaccine’s ability to establish a population of pulmonary tissue resident memory (T_RM_) T cells since these cells have been shown to provide a first line of defense against invading pathogens (11).

## MATERIALS AND METHODS

### Mice

Male C57BL/6 mice were obtained from the Jackson Laboratories (Bar Harbour, Maine). Mice used in this study were 6 – 8 weeks old and housed at the Center for Biologics Evaluation and Research, Silver Spring, MD in a barrier environment. This study was done in accordance with ARRIVE guidelines for the care and use of laboratory animals specified by the National Institutes of Health and was approved by the Institutional Animal Care and Use Committee of the Center for Biologics Evaluation and Research under Animal Study Protocol 1993-09 (12).

### Immunizations

BCG vaccine (Pasteur strain) for all BCG containing vaccine formulations was administered subcutaneously (SC) in PBS at 1 × 10^6^ CFU per immunization unless otherwise stated. The adjuvant-containing BCG formulation was prepared by mixing BCG with dimethyl dioctadecyl-ammonium bromide (1.5 mg/ml final in water) (DDA, Kodak, Rochester, NY) and D-(+)-Trehalose 6,6’-Dibehenate (0.5 mg/ml final in water) (TDB, Avanti Polar Lipids, Alabaster, AL) as previously described (7). For the BAA vaccine, BCG formulated in the above adjuvant was prepared with rEsat-6 (25 µg/ml final) obtained from BEI Resources (NR-49424) (Manassas, VA) or purified in-house from *Escherichia coli* cultures expressing the recombinant protein as described previously (13). We chose the Esat-6 antigen since this is an immunodominant *M. tuberculosis* antigen (15). We administered BCG, the adjuvant alone or BCG+Adj with or without rEsat-6 formulations three times, two weeks apart subcutaneously (SC) in a volume of 0.2 ml. The adenovirus construct expressing an Esat-6, Antigen 85B gene fusion (AdE6-85B) was created as previously described and produced by Viraquest (North Liberty, IA) (16). The AdE6-85B construct was administered intranasally (IN) 6 weeks after the final BCG+Adj/rEsat-6 immunization by delivering 5 × 10^8^ pfu into the nares of anesthetized mice (ketamine + xylazine cocktail administered intraperitoneally) via pipette tip in a volume of 25 µl. We designated the BCG+Adj/rEsat-6 prime, AdE6-85B boost vaccine regimen as BAA.

### Latency model

For evaluating vaccine efficacy via the mouse latency model, the animals were immunized with different vaccine formulations or routes to discover the most protective vaccine or vaccine regimen or route of administration. Specifically, we tested BCG administered intravenously (IV) in the tail vein or formulated in adjuvant with or without rEsat-6 (BCG+Adj or BCG+Adj/rEsat-6) with or without a AdE6-85B IN boost. Eight weeks after the final SC immunization, the mice were challenged via aerosol with around 100 CFU *M. tuberculosis* Erdman strain using a Middlebrook chamber (Glas Col, Terre Haute, IN). One month post-challenge, the mice were administered isoniazid (200 mg/Liter) (INH) and rifampin (100 mg/Liter) (Rif) (INH/Rif water) orally in their drinking water ad libitum for three months to reduce the pulmonary infection to undetectable levels without clearing the infection. Five mice were euthanized at the end of the INH/Rif treatment to verify the absence of detectable CFU in the lungs, or we waited three months post-INH/Rif to evaluate vaccine efficacy in the remainder of the mice. To assess pulmonary mycobacterial burdens, we plated 0.5 ml of a 1:5 dilution of the lung homogenates (5 ml volume in PBS with 0.05% Tween-80 using a Seward Stomacher 80 Blender) (Tekmar, Cincinnati, OH) from separate mice (4 – 10 mice per group) per 7H11 agar plate supplemented with 10% OADC enrichment (Becton Dickinson, Sparks, MD), 10 μg/ml ampicillin, 50 μg/ml cycloheximide, and 2 µg/ml 2-thiophenecarboxylic acid hydride (TCH) (Sigma, St. Louis, MO) (7). The concentration of TCH added to the agar plates inhibits BCG growth while permitting *M. tuberculosis* growth. Plates were incubated at 37°C for 4 weeks before counting to determine the number of mycobacterial colony-forming units (CFU) in the lungs. We considered the presence of only one colony on a plate as indicating a mouse had progressed to active disease. A timeline illustrating the latency model is provided in Fig. 1A.

**FIG 1.**
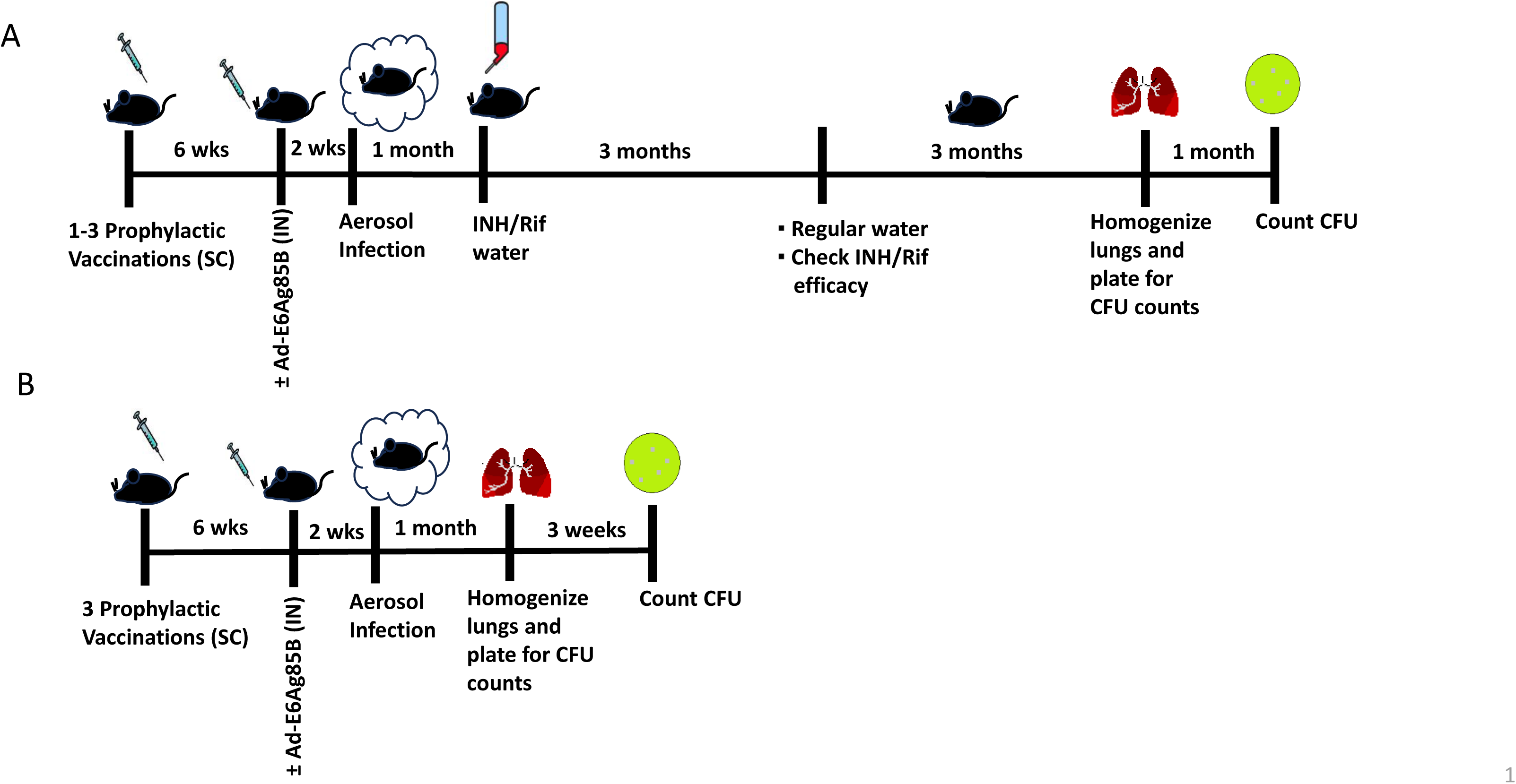
Timelines illustrating the mouse latency and acute infection models. (**A**) The latency model was performed as follows: six weeks following the final SC prophylactic immunization, only mice vaccinated with BCG+Adj/Esat-6 or adjuvant alone were immunized IN with the adenovirus construct expressing an Esat-6-Ag85B fusion (Ad-E6Ag85B) (BAA). The mice then received an aerosol challenge with ∼100 CFU *M. tuberculosis* Erdman. One month post-challenge the mice were administered drinking water (ad libitum) containing INH and Rif to reduce the pulmonary mycobacterial burden to below detectable levels without clearing the infection. Following three months of INH/Rif water treatment, the mice per placed on regular drinking water. At this time, five mice were euthanized to verify that there were no detectable CFU in the lungs. Three months following cessation of INH/Rif treatment the mice were euthanized, their lungs were homogenized, and aliquots of the homogenates were plated on 7H11/OADC plates. After a one month incubation time, CFU growing on the plates were counted to assess the ability of the vaccines to prevent active disease (N = 4 – 10 mice per group). (**B**) The acute infection model was performed as described for the latency model up to the time of INH/Rif treatment (1 month post-challenge). Instead of the INH/Rif treatment, the mice were euthanized, the lungs were homogenized, and aliquots were plated on 7H11/OADC plates to assess the ability of the vaccine induced immune response to control mycobacterial growth in the lungs (N = 4 – 5 mice per group).

### Acute infection model

Eight weeks after the final SC immunization with BCG, adjuvant alone, BCG/Adj or BAA (2 weeks post-AdE6-85B IN boost), 4 - 5 mice per group were infected with around 100 CFU *M. tuberculosis* (Erdman strain) by aerosol in a Middlebrook chamber. Two or four weeks post-challenge, the lungs were homogenized separately as described below for lung cell isolation for flow cytometry analysis. Homogenate aliquots were serially diluted in PBS + 0.05% Tween-80 and plated on Middlebrook 7H11/OADC agar (Difco) plates containing 10% OADC enrichment with antibiotics (10 μg/ml ampicillin, 50 μg/ml cycloheximide, and 2 µg/ml TCH) (7). Plates were incubated at 37°C for 3 weeks, and the number of mycobacterial CFU in the lungs was determined using a colony counter. Protection was defined as the organ log_10_ CFU difference between nonimmunized (Naïve) and immunized mice. The acute infection model is illustrated in Fig. 1B.

### Flow Cytometry

Pulmonary cells were characterized by flow cytometry and antibody staining to determine the frequency of T cells expressing cytokines or surface markers. Lung cells were obtained either before challenge or two or four weeks following an aerosol challenge with *M. tuberculosis* Erdman by disrupting the lungs with razor blades and incubating the tissue with Dispase (2 Units/ml) (Corning, Glendale, AZ) in PBS + 2% FBS (PBS/FBS) for 1 hour at 37°C. The lung homogenates were then passed through filter bags (Whirl-Pak, Pleasant Prairie, WI) to remove tissue clumps and then treated with ACK lysing buffer (Lonza, Walkersville, MD) for 5 min. at room temperature (RT) to remove erythrocytes. The cells were then passed through 70 µm cell strainers to remove cell debris. Typically, 3 – 10 × 10^6^ cells were recovered from the lungs of individual mice. After washing the cells with PBS containing 0.5% FBS and 2 mM EDTA (FACS buffer), the cells were incubated with Brefeldin A in DMEM supplemented with 10% FBS, 10 mM HEPES, 2.0 mM L-glutamine, and 0.1 mM minimal essential medium nonessential amino acids for 4 hours in 6-well plates (3 ml per well). The cells were then harvested from the plates and transferred to separate 12 × 75 mm tubes, pelleted (400 x g for 5 min.) and resuspended in ≤ 0.1 ml volume and then incubated with Near-IR (NIR) live-dead stain (10 µl of a 1:100 dilution) (Invitrogen) for 15 min. at 4°C to allow gating on viable cells. Antibody against CD16/CD32 (FcγIII/II receptor, clone 2.4G2) (Fc block) was also added and incubated at 4°C for 15 min.

Surface staining was accomplished by adding fluorochrome-conjugated antibodies specific for the following molecules and then incubated at 4°C for 30 min.: TCRβ (AF700; clone H57-597), CD4 (PerCP-Cy5.5; clone GK1.5), CD8 (BV421; clone 53-6.7), CD153 (PE; clone RM153), CXCR3 (PE-Cy7, Clone CXCR3-173), PD-1 (PE, clone 29F.1A12), CD69 (BV421, clone H1:2F3), P2X7R (APC, clone 1F11) or KLRG1 (FITC, clone 2F1) (0.1 – 0.2 µg each). Following surface staining, the cells were washed with 2.0 ml FACS buffer and then fixed with 2% paraformaldehyde (in PBS) for 30 min. and then washed with PBS/FBS. Detection of intracellular IFN-γ (APC, clone XMG1.2), TGFβ (BV421, clone TW7-16B4) and/or IL-17A (PE, clone TC11-18H10) was done as previously described (7). Briefly, the fixed cells were washed and permeabilized with 2.0 ml perm/wash buffer (0.1% saponin, 10 mM Hepes, 1% FBS) and then resuspended in ≤ 0.1 ml perm/wash buffer followed by the addition of antibodies (0.2 µg each). After a 45 min. incubation at room temperature in the dark the cells were washed once with 2.0 ml perm/wash buffer followed by a wash with 2.0 ml PBS/FBS. Fluorescence minus one (FMO) controls were included to facilitate gating the positively stained cell populations (gating scheme and representative FMO control dot plot panels are provided in Fig. S1) The antibodies were purchased from BD Biosciences or Biolegend. The fixed cells were analyzed using a LSR Fortessa flow cytometer (Becton Dickinson) and FlowJo software (Tree Star Inc., Ashland, OR) collecting 100,000 to 200,000 events.

### Statistical analysis

Graph Pad Prism 5 software was used to analyze the data for these experiments (Graph Pad Software, San Diego CA). Protection and flow cytometry results were evaluated using two-tailed, unpaired Mann-Whitney t-test analysis when comparing two experimental groups or one-way ANOVA with Tukey’s post-test for multi-group comparisons. Protection and flow cytometer results are expressed as the mean ± the standard deviation (SD).

## RESULTS

We utilized the mouse *M. tuberculosis* latency model described in the Methods section and illustrated in Fig. 1A to screen vaccines that reduced the frequency of reactivation in a majority of immunized animals and relative to nonimmunized mice. In order to characterize vaccine-induced immune responses of only highly effective vaccines we considered the presence of at least one colony on the 7H11 agar plates inoculated with aliquots of infected lung homogenates as indicative of a mouse having reactivated (progressed to active disease). For these experiments, we verified the absence of detectable CFU in the lungs from five mice at the end of the INH/Rif treatment and then assessed the mycobacterial burdens in mice from each group three months later. Table 1 contains reactivation frequency data using the latency model and shows 90 – 100% of nonimmunized control mice consistently progressed to active disease three months following cessation of INH/Rif treatment which explains why we chose the three month post-INH/Rif treatment time point to evaluate vaccine efficacy. In contrast, the BAA vaccine consistently protected 75 – 100% of immunized mice against reactivation. The other vaccines were either nonprotective (e.g. BCG administered intravenously) or demonstrated varying degrees of efficacy and were not as consistently protective as the BAA vaccine (e.g. BCG+Adj or BCG+Adj/rEsat-6). For this reason, the focus of subsequent experiments was the BAA vaccine formulation. Interestingly, among the immunized mice that did reactivate, significant differences in pulmonary CFU counts relative to nonimmunized control mice were only observed for mice immunized with vaccines that prevented reactivation in a majority of the immunized animals (BCG+Adj, BCG+Adj/Esat-6 and BAA) (p < 0.02).

**TABLE 1.**
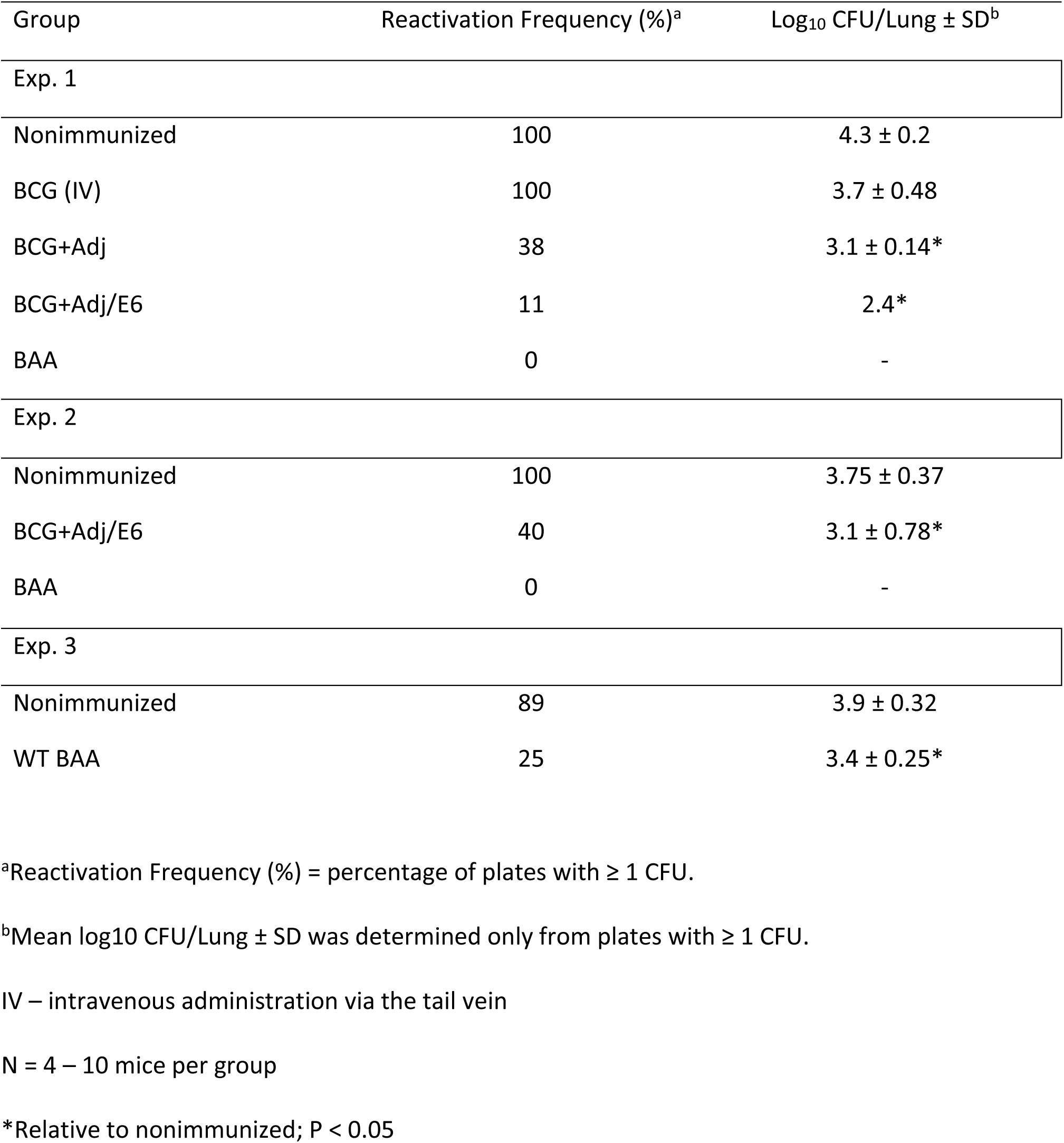
Protection using the latency model.

| Group | Reactivation Frequency (%)<sup>a</sup> | Log<sub>10</sub> CFU/Lung ± SD<sup>b</sup> |
| --- | --- | --- |
| Exp. 1 |  |  |
| Nonimmunized | 100 | 4.3 ± 0.2 |
| BCG (IV) | 100 | 3.7 ± 0.48 |
| BCG+Adj | 38 | 3.1 ± 0.14* |
| BCG+Adj/E6 | 11 | 2.4* |
| BAA | 0 | - |
| Exp. 2 |  |  |
| Nonimmunized | 100 | 3.75 ± 0.37 |
| BCG+Adj/E6 | 40 | 3.1 ± 0.78* |
| BAA | 0 | - |
| Exp. 3 |  |  |
| Nonimmunized | 89 | 3.9 ± 0.32 |
| WT BAA | 25 | 3.4 ± 0.25* |

<sup>a</sup>Reactivation Frequency (%) = percentage of plates with ≥ 1 CFU.
<sup>b</sup>Mean log<sub>10</sub> CFU/Lung ± SD was determined only from plates with ≥ 1 CFU.
IV – intravenous administration via the tail vein
N = 4 – 10 mice per group
\*Relative to nonimmunized; P < 0.05

We next evaluated the efficacy of the BAA vaccine using an acute infection mouse model as illustrated in Fig. 1B and described in the Methods section. Either three weeks or one month following an aerosol *M. tuberculosis* challenge, mice immunized with the BAA vaccine limited a pulmonary infection significantly better than control groups (Naïve, Adj+Ad, BCG, or BCG+Adj) (p < 0.05) with an impressive 2.5 – 3 log_10_ CFU reduction relative to Naïve mice (p < 0.0001) (Fig. 2A, B). The other vaccines (excepting Adj + Ad) were also significantly protective relative to Naïve mice (p < 0.0001).

**FIG 2.**
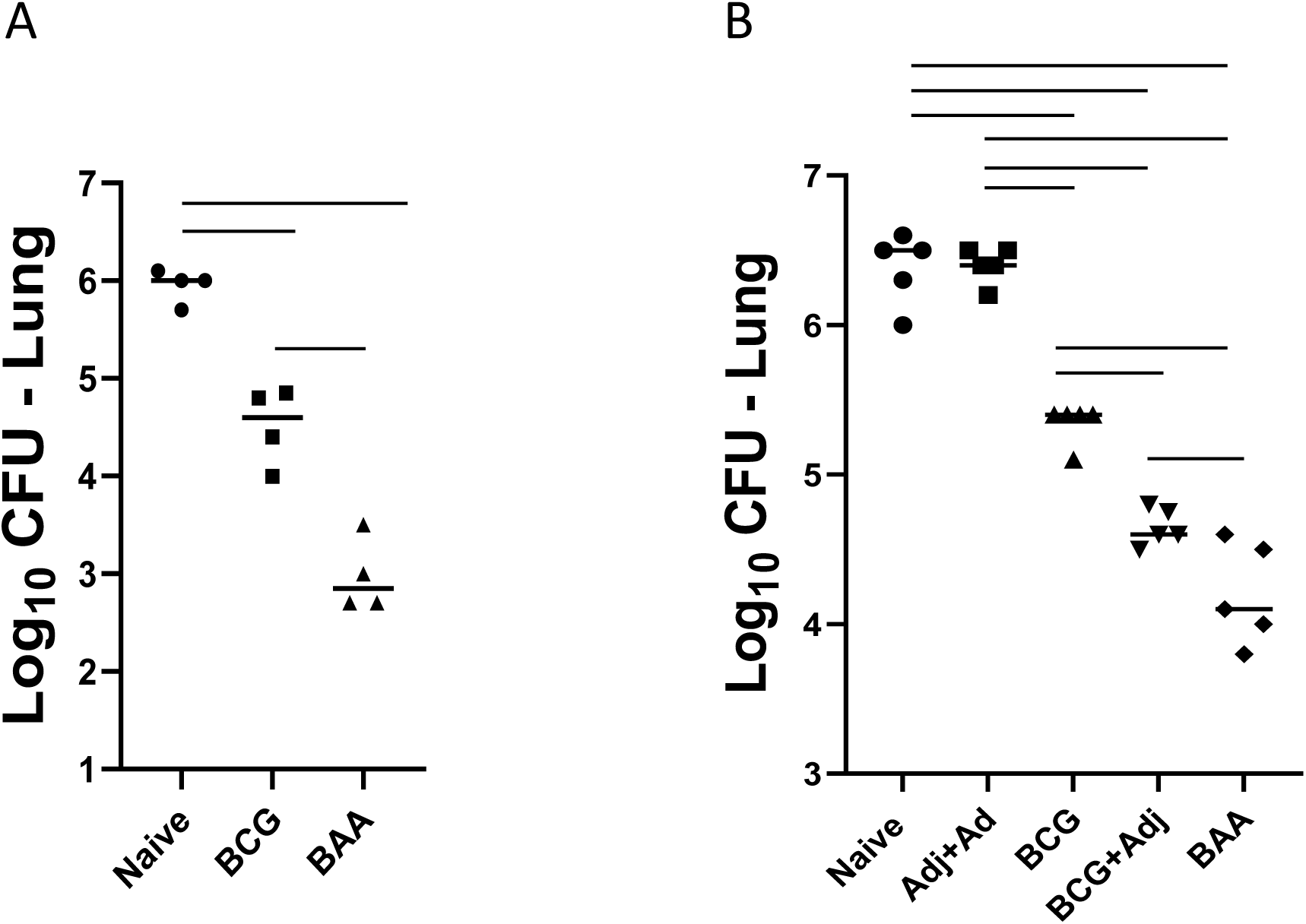
Protective efficacy of the BAA vaccine is significantly augmented relative to control vaccines using the acute infection model. Mice were vaccinated SC three times, 2 weeks apart with either BCG, BCG+Adj, Adjuvant alone with an AdE6-85B IN boost (Adj + Ad) or BCG+Adj/Esat-6 with the AdE6-85B IN boost 6 weeks after the third SC immunization (BAA). Eight weeks after the final SC immunization or two weeks post-AdE6-85B IN immunization, the mice received an aerosol dose of ∼100 CFU *M. tuberculosis* Erdman. Mean (± SD) pulmonary CFU were determined from the lungs at either three (**A**) or four (**B**) weeks post-challenge. Horizontal bars indicate statistical significance between the different groups (p < 0.05). N = 4 – 5 mice per group.

Using flow cytometry, we interrogated pulmonary cells from the different groups using antibodies against known immune markers and cytokines before challenge and at two weeks following an *M. tuberculosis* aerosol infection. Since CD153, a member of the TNF superfamily, was previously shown to be a marker for protective CD4^+^ T cells (17–19) we examined the lungs of BAA immunized animals and control groups for expression of this molecule. As shown in Fig. 3A, pre-challenge frequencies of CD4^+^ T cells expressing CD153 were significantly elevated in the lungs of animals immunized with BAA relative to the control groups (p < 0.0001). In a separate experiment, the number of CD4^+^CD153^+^ T cells recovered from the lungs of BAA vaccinated mice was significantly greater than the number recovered from Naïve or BCG groups (3,819 ± 1711, 296 ± 121 and 425 ± 202 respectively; p < 0.0005) (not shown). At two weeks post-challenge, the number of CD4^+^ T cells expressing CD153 was only significantly elevated in the lungs of the BCG+Adj and BAA groups relative to Naïve mice and the other control groups (p < 0.05) (Fig. 3B). Representative dot plots showing pre-challenge frequencies of CD4^+^CD153^+^ cells from the different groups are shown in Fig. 3C.

**FIG 3.**
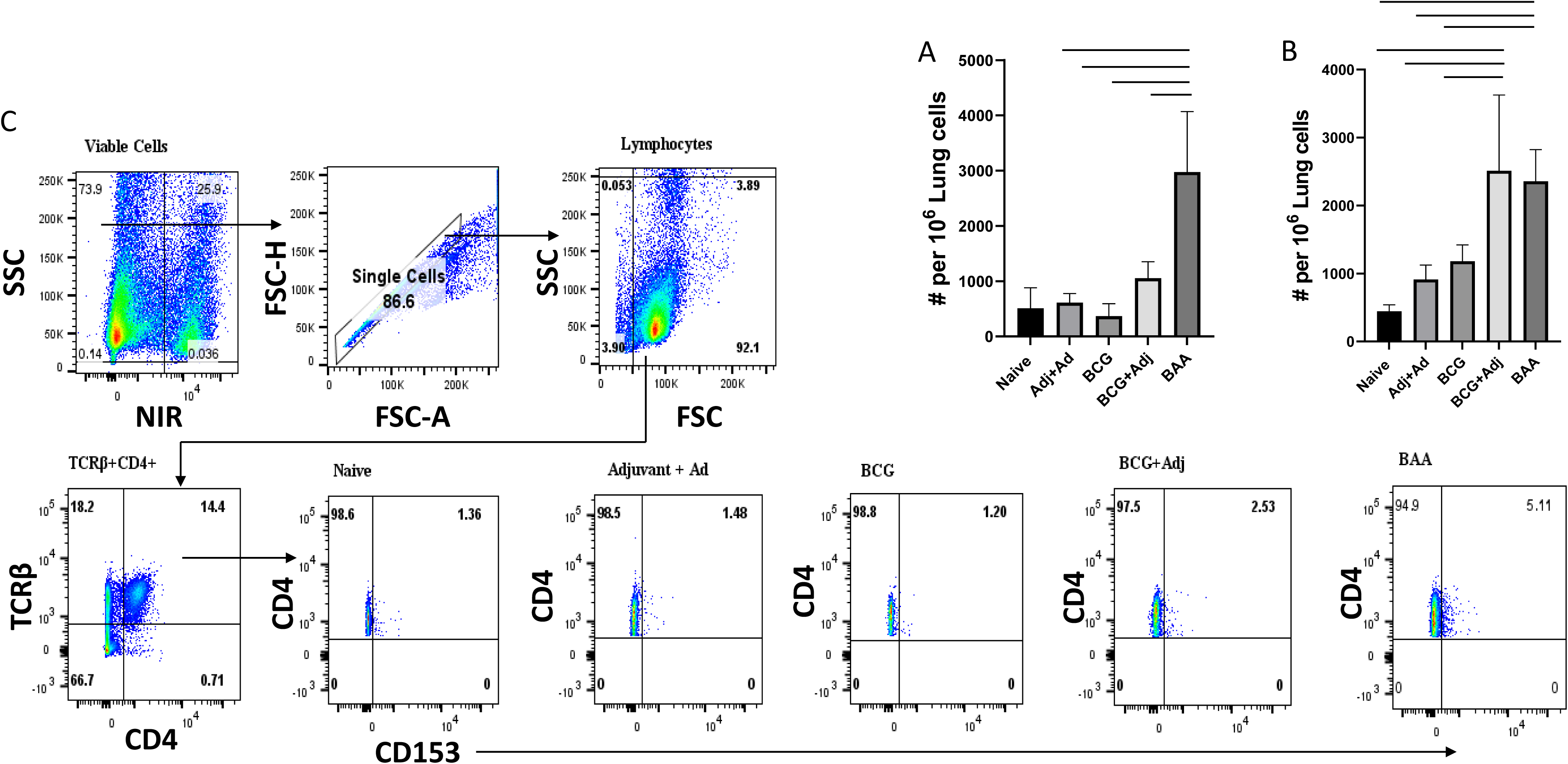
The frequency of pulmonary TCRβ^+^CD4^+^ cells expressing the CD153 molecule is significantly elevated in mice immunized with BAA before or after challenge. Mice were vaccinated with Adj + Ad, BCG, BCG + Adj or BAA as described above in the legend for Fig. 1 and then, using flow cytometry, the pulmonary TCRβ^+^CD4^+^ cells were examined for expression of the CD153 marker. The number of TCRβ^+^CD4^+^ cells per million lung cells expressing CD153 is shown for the different groups (Naïve, Adj + Ad, BCG, BCG + Adj or BAA) either before (**A**) or following (**B**) an *M. tuberculosis* Erdman aerosol challenge (∼100 CFU). Representative dot plot (**C**) showing the frequencies (%) of TCRβ^+^CD4^+^ cells expressing CD153. Horizontal bars indicate statistical significance between the different groups (p < 0.05). N = 4-5 mice per group.

Expression of the P2X7R molecule by CD4^+^ T cells has also been shown to be associated with control of mycobacterial growth in the lungs (20, 21). Although we did not find post-challenge elevated expression of P2X7R by pulmonary CD4^+^ T cells from the BAA group relative to the other groups, we did see significant differences prior to challenge (Fig. 4B) (p < 0.02). Only the BAA vaccine regimen induced significantly elevated frequencies of CD4^+^P2X7R^+^ T cells relative to all the other groups with no significant differences between the other groups. In a separate experiment, we also observed significantly elevated frequencies of CD4^+^P2X7R^+^ T cells in the lungs before challenge from mice immunized with the BAA vaccine relative to BCG immunized mice and Naïve mice (15,458 ± 6782, 1,450 ± 775 and 811 ± 501 respectively; p < 0.001) (not shown). Only a very small percentage of these cells expressed the KLRG1 molecule indicating that a majority of these cells were most likely protective, residing in the parenchyma rather than the lung vasculature (Fig. 4A).

**FIG 4.**
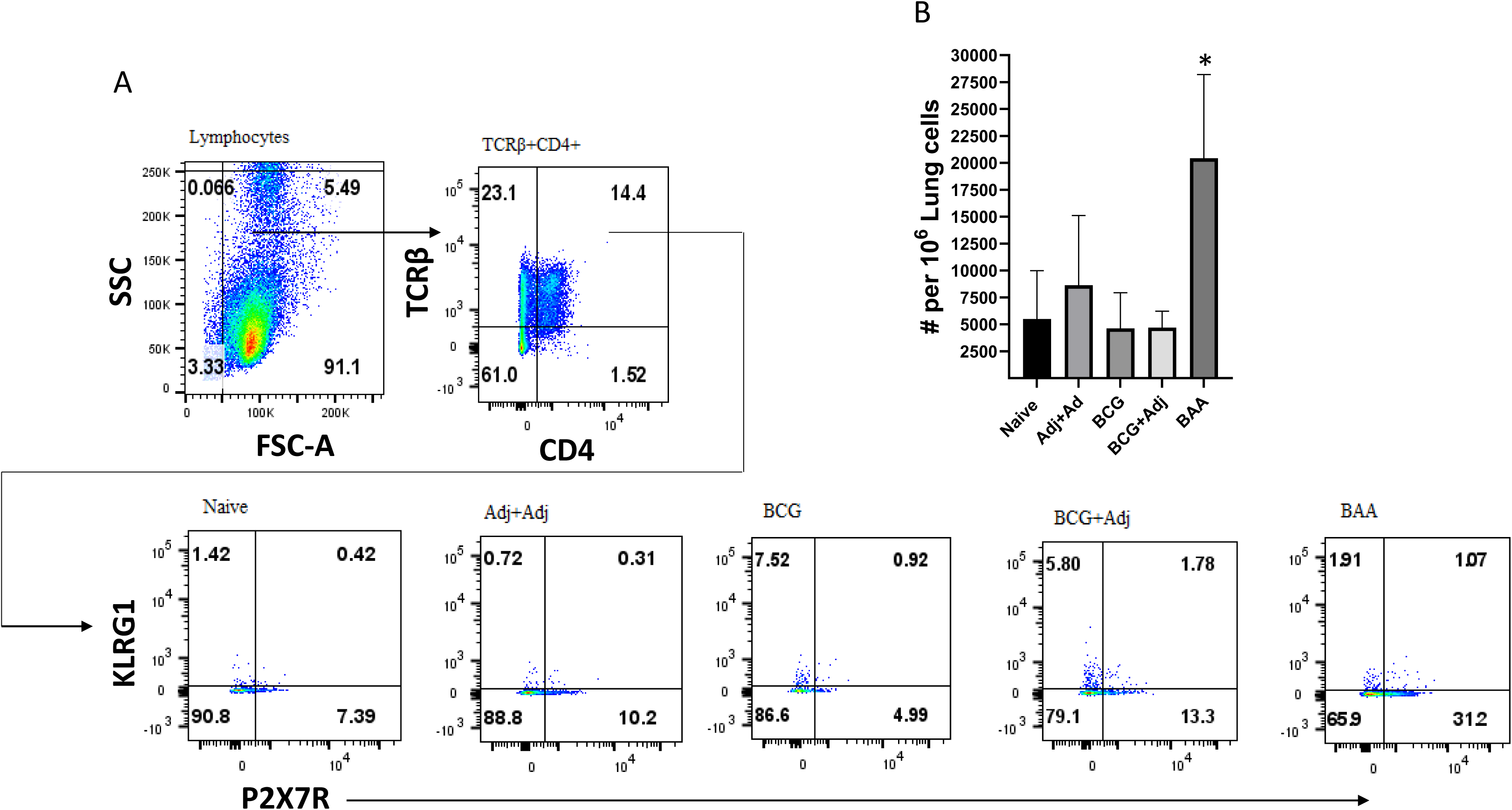
The frequency of pulmonary TCRβ^+^CD4^+^ cells expressing the P2X7R molecule is significantly elevated in mice immunized with BAA before challenge. Mice were vaccinated with Adj + Ad, BCG, BCG + Adj or BAA as described above in the legend for Fig. 1 and then, using flow cytometry, the pulmonary TCRβ^+^CD4^+^ cells were examined for expression of the P2X7R molecule. Representative dot plot (**A**) showing the frequencies (%) of TCRβ^+^CD4^+^ cells expressing P2X7R. (**B**) The number of TCRβ^+^CD4^+^ cells per million lung cells expressing P2X7R are shown for the different groups (Naïve, Adj + Ad, BCG, BCG + Adj or BAA) prior to challenge. (*p < 0.05). N = 4-5 mice per group.

Since tissue resident memory (T_RM_) T cells have been shown to be involved in protection against *M. tuberculosis*, we wanted to examine the frequencies of these cells post-vaccination both before and after challenge in the lungs of the different groups (22). These cells have been defined in previous studies as expressing CD69, PD-1 and CXCR3 molecules (22–25). We found mostly CD4^+^ T cells expressing these molecules in the lungs with very few CD8^+^ T_RM_ cells detected (not shown). We observed the highest frequencies of these cells in the lungs of BAA immunized mice before challenge which were significantly elevated relative to control groups (p < 0.05) (Fig. 5A). Interestingly, only T_RM_ frequencies in the Adj + Ad and BAA groups were significantly elevated relative to the Naïve group (p < 0.02 and 0.0001 respectively). Representative dot plots showing pre-challenge frequencies of T_RM_ cells in the lungs of mice immunized with the BAA vaccine are shown in Fig. 5C. In a separate experiment we again found that frequencies of T_RM_ cells in the lungs of BAA vaccinated mice were significantly elevated relative to Naïve and BCG immunized mice before challenge (1151 ± 730, 39 ± 32 and 43 ± 16 T_RM_ cells per million lung cells respectively) (p < 0.01) (not shown). Remarkably, by two weeks post-challenge, very few T_RM_ cells were detected in the lungs of any of the groups indicating that these cells are either rapidly depleted, lose one or more identifying markers or convert to another T cell type after challenge (not shown). Of note, the great majority of T_RM_ CD4^+^ T cells do not express KLRG1 which is noteworthy since expression of this molecule has been shown to identify a population of terminally differentiated, vascular pulmonary CD4^+^ T cells that do not contribute to protection (Fig. 5C) (19, 23, 25). The number of pulmonary CD4^+^ T_RM_ cells expressing the P2X7R molecule was significantly higher in the BAA group relative to the other groups (p < 0.002) (Fig. 5B). In a separate experiment, frequencies of CD4^+^ T_RM_ P2X7R^+^ cells were similarly significantly elevated in the lungs of mice immunized with BAA relative to Naïve and BCG vaccinated controls before challenge (498 ± 319, 76 ± 75, 64 ± 34 per million cells respectively) (p < 0.02) (not shown).

**FIG 5.**
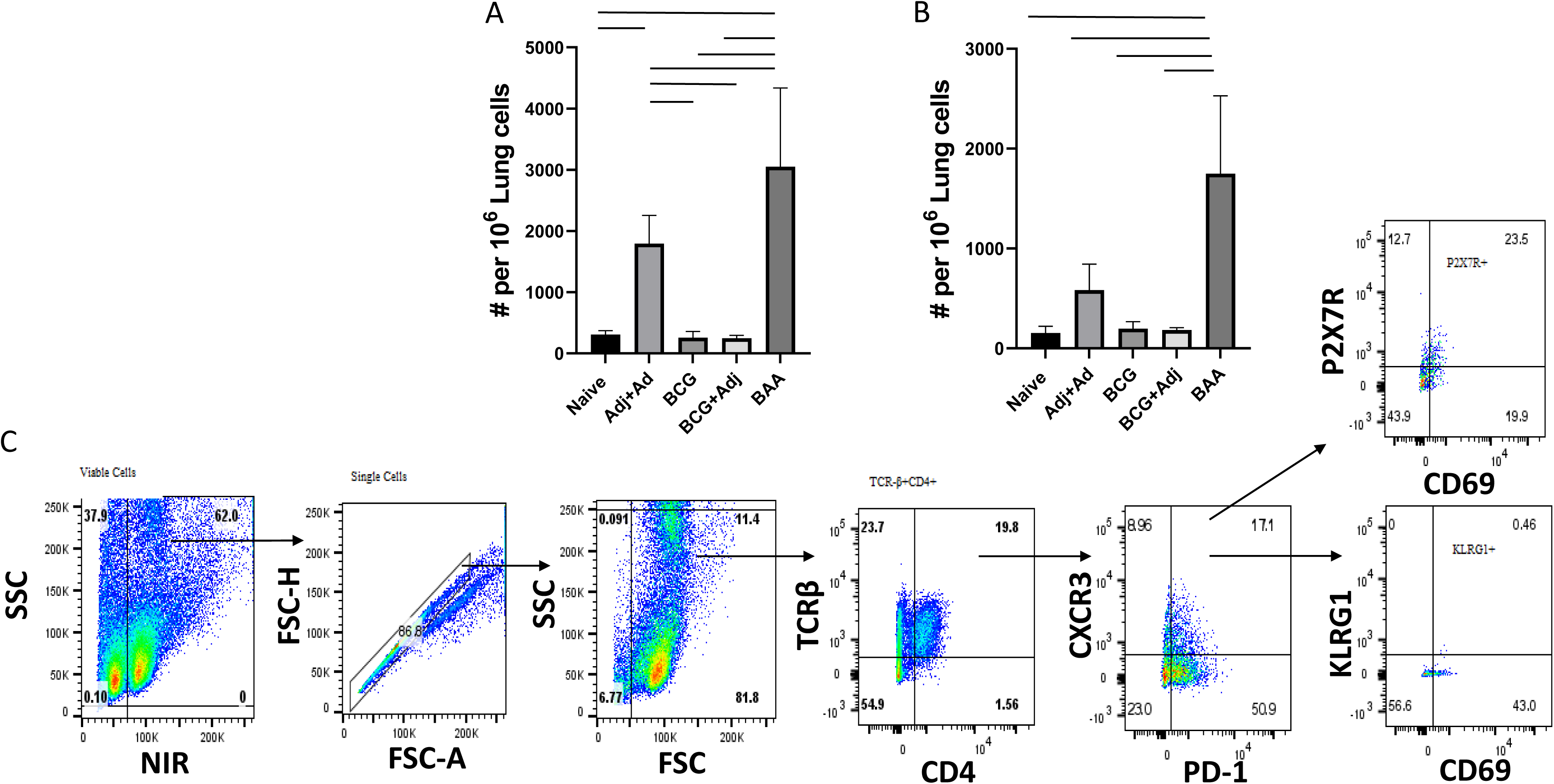
The pre-challenge frequency of pulmonary TCRβ^+^CD4^+^ Tissue Resident Memory (T_RM_) cells is significantly elevated in mice immunized with BAA. (**A**) The number of pre-challenge CD4^+^ Tissue Resident Memory (PD-1^+^CD69^+^CXCR3^+^) T cells or (**B**) CD4^+^ T_RM_ P2X7R^+^ T cells per million lung cells was assessed in the lungs of Naïve or vaccinated mice (Adj + Ad, BCG, BCG + Adj or BAA) by flow cytometry. (**C**) Representative dot plot showing the gating scheme for assessing the frequencies (%) of pulmonary TCRβ^+^CD4^+^ T_RM_ cells. Horizontal bars indicate statistical significance between the different groups (p < 0.05). N = 4 – 5 mice per group.

We also examined expression of different cytokines after challenge in the lungs of the different groups. In addition to IFN-γ and IL-17A, which have been shown to be important for immunity against *M. tuberculosis*, we also examined expression of TGFβ since this cytokine has been shown to be involved in the induction and maintenance of T_RM_ cells at mucosal sites of infection (28, 29). We found that CD4^+^TGFβ^+^ frequencies were only significantly elevated in the lungs of mice immunized with the BAA vaccine relative to the other groups but only after challenge (Fig. 6) (p < 0.001). At one month post-challenge, frequencies of CD4^+^ T cells expressing only IL-17A (CD4^+^IL-17^+^) were significantly elevated in the lungs of mice immunized with BCG+Adj and BAA relative to the other groups (p < 0.01), and the frequency of CD4^+^ T cells expressing either TGFβ and IL-17 or TGFβ, IL-17 and IFN-γ were also significantly elevated in the lungs of only the BAA group (p < 0.05). Of note, we also observed significant differences in the frequencies of CD4^+^ cells expressing only IL-17^+^ in the lungs of BCG+Adj relative to the BCG, Adj+Ad and Naïve groups (p < 0.01). Surprisingly, the frequencies of cells expressing only IFN-γ were significantly elevated in the lungs of Naïve mice and mice immunized with Adj + Ad relative to the protected BCG+Adj and BAA groups (p < 0.05). Representative dot plots showing gating for determining the frequencies of monofunctional and multifunctional cell populations from the lungs of Naïve and BAA immunized mice are shown in Fig. S2.

**FIG 6.**
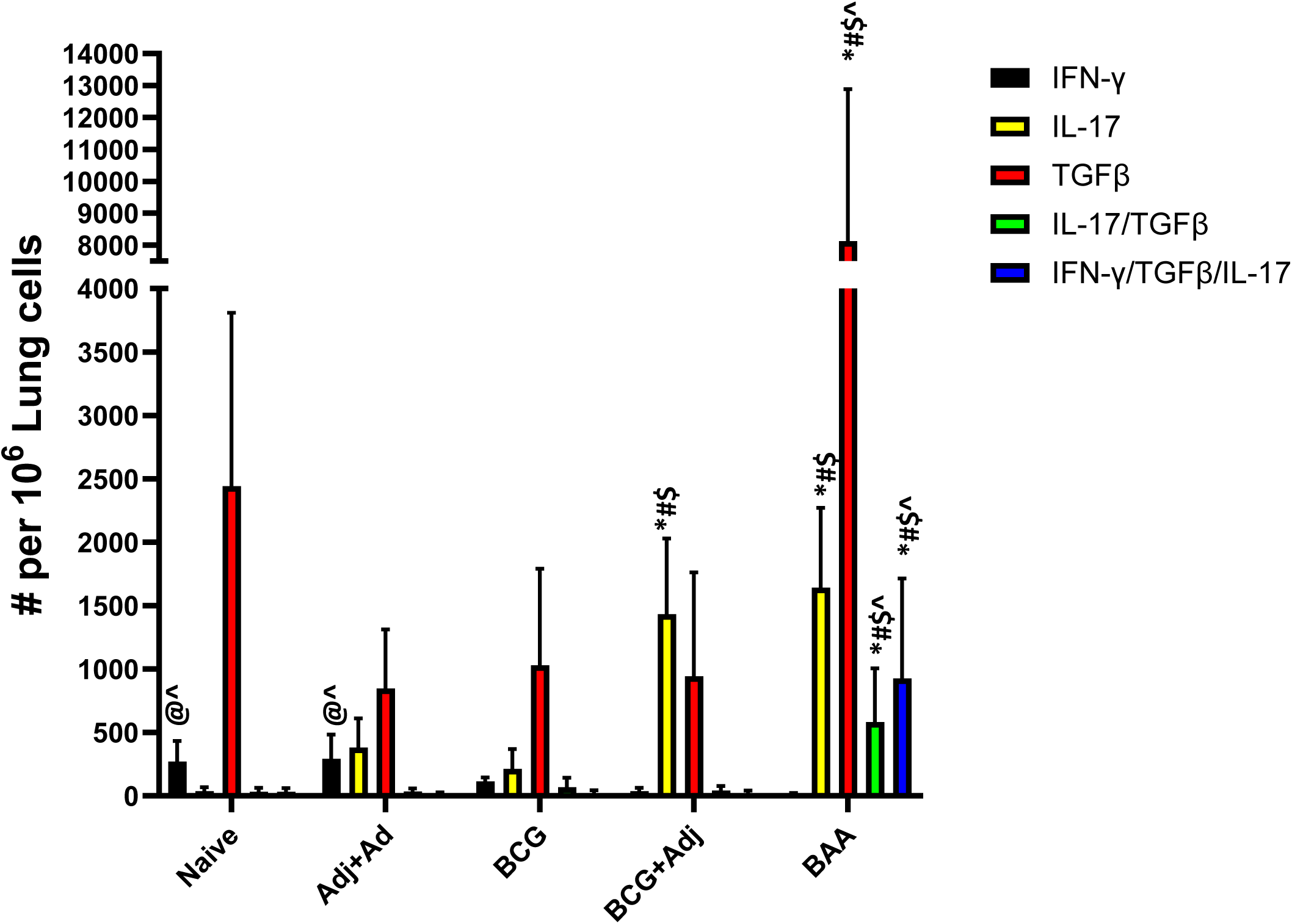
The post-challenge frequency of pulmonary multifunctional TCRβ^+^CD4^+^ cells expressing one or more cytokines was assessed from the different groups. At one month post-challenge, the number of pulmonary CD4^+^ T cells per million lung cells from the different groups (Naïve, Adj + Ad, BCG, BCG + Adj or BAA) was assessed by flow cytometry for expression of only IFN-γ, IL-17 or TGFβ or a combination of two cytokines (IFN-γ^+^IL-17^+^) or all three cytokines (IFN-γ^+^TGFβ^+^IL-17A^+^). Statistical significance relative to Naïve (*), Adj+Ad (^#^), BCG (^$^), BCG+Adj (^^^) or BAA (^@^) is indicated above the vertical graph bars ( *P* < 0.05). N = 4 – 5 mice per group.

## DISCUSSION

We chose to utilize a mouse TB latency model to screen potential TB vaccine candidates since this is a more stringent model than the acute infection model and, ideally, a new TB vaccine should reduce reactivation rates in order to interrupt transmission. Moreover, our ultimate objective of this study was to characterize pulmonary immunity induced by vaccines that consistently reduce reactivation rates. Importantly, the INH/Rif water treatment consistently reduced the pulmonary infection to undetectable levels in all mice examined, and since virtually all the nonimmunized mice consistently progressed to active disease, the INH/Rif treatment regimen did not clear the infection. Our initial work using the latency model found three vaccines that consistently reduced reactivation rates: BCG+Adj, BCG+Adj/E6 and BAA; BAA, however, was the most consistent. The BAA vaccine consistently protected 75 – 100% of immunized animals from progressing to active disease over a three month period; in contrast, 90 – 100% of nonimmunized control mice consistently progressed to active disease over the same time. These three vaccines were also remarkable for their ability to control mycobacterial growth in the lungs significantly better than nonimmunized controls. Of note, BCG delivered IV was unable to reduce reactivation rates or reduce pulmonary bacterial burdens relative to nonimmunized control mice indicating that BCG formulated in DDA/TDB adjuvant was necessary for any protection using this model.

The BAA vaccine was also highly and significantly protective relative to Naïve controls and other vaccines tested (Adjuvant alone, BCG, BCG+Adj) using the acute infection model. Additionally, we examined the pulmonary immune response in Naïve and immunized mice both before and following an aerosol *M. tuberculosis* challenge to compare mice immunized with BAA to the other groups and observed some important differences. Since expression of CD153 by T cells has been shown to be necessary for control of mycobacterial growth in mice, and since CD4^+^CD153^+^ T cell frequencies have been reported in the BAL and granulomas of infected monkeys and were inversely correlated with bacterial burden (17, 19), we examined CD153 expression from T cells in the lungs of the different groups using flow cytometry. We found that the number of pulmonary CD4^+^ T cells expressing this marker from mice immunized with BAA was significantly increased relative to the other groups before challenge and was also significantly elevated two weeks post-challenge in both the BAA and BCG+Adj groups relative to the other groups. This finding is noteworthy since CD153 is a T cell co-stimulating molecule signaling T cell proliferation and secretion of effector molecules, and CD153 KO mice were previously shown to have a reduced ability to control mycobacterial infections relative to wild-type (wt) mice (27, 28). Others have shown that expression of CD153 by CD4^+^ T cells was significantly reduced from patients with active disease versus those with a latent infection (19), and the CD4^+^CD153^+^ T cell response inversely correlated with sputum bacterial CFU (18). Our BAA protection results using both mouse models and flow cytometry results are consistent with these previous findings in animal models and in humans and suggest that vaccine-mediated expansion of pulmonary CD4^+^CD153^+^ T cells may be predictive of greater control of an acute mycobacterial infection and may serve as a marker of protection against progression to active disease.

Tissue resident memory (T_RM_) cells serve as a first line of defense against invading pathogens at different mucosal sites. Since the presence of CD4^+^ T_RM_ cells in the lungs has been shown to contribute to immunity to TB, we were interested in comparing the frequencies of these cells in the lungs of the different groups (22, 25). T cells expressing the CD69, CXCR3 and PD-1 molecules were previously identified as markers for CD4^+^ T_RM_ T cells (22–24), and we found that frequencies of pulmonary CD4^+^ T cells expressing these markers were significantly elevated in the BAA group relative to the other groups. We also observed that relatively few of these cells express the KLRG1 molecule (by any group) which is a marker for vasculature, nonprotective, terminally differentiated CD4^+^ T cells which do not migrate into the lung parenchyma (19, 25). In contrast, a majority of these cells expressed the P2X7R marker which is a proinflammatory, purinergic receptor binding to extracellular ATP or NAD, which become elevated during pathogen-mediated tissue damage. While lower concentrations of ATP or NAD at the site of tissue damage (i.e. infection) leads to T cell activation and proliferation, prolonged activation by ATP or NAD leads to T cell death via formation of P2X7R pores which may, at least, partially explain why we were unable to detect pulmonary T_RM_ cells one month following an *M. tuberculosis* challenge (30, 31). Importantly, CD38, TGFβ and extracellular nucleotide signaling via P2X7R were previously found to be involved in vaccine induced development and maintenance of CD4^+^ T_RM_ cells in the gastric epithelium in a mouse model for *Helicobacter pylori* infection (26). Interestingly, T_RM_ frequencies in the Adj + Ad group were also significantly elevated relative to all other groups except for BAA which may indicate that the Adenovirus construct delivered IN is mostly responsible for the establishment of T_RM_ cells in the lungs. However, given that the Adj + Ad vaccine is not protective, qualitative differences likely exist in the population of T_RM_ cells induced by these two vaccines in addition to differences in P2X7R expression.

Interestingly, it has been shown that CD30/CD153 signaling promotes differentiation of Th17 cells (32), and we previously demonstrated that IL-17A expression was associated with superior protection mediated by the BCG+Adj vaccine relative to BCG alone (10). We also found in the present study that frequencies of CD4^+^ T cells expressing IL-17A alone were significantly elevated in the lungs of both BAA and BCG+Adj immunized mice one month post-challenge relative to the other groups; however, significantly elevated frequencies of CD4^+^ cells expressing both IL-17 and TGFβ or TGFβ alone or expressing IFN-γ, IL-17 and TGFβ were only significantly elevated in the BAA group relative to the other groups. We examined expression of TGFβ since, as mentioned above, this cytokine has been shown to promote differentiation of CD4^+^ T_RM_ cells (26). Unexpectedly, frequencies of cells expressing IFN-γ alone were observed only in the nonprotected Naïve and Adj + Adj groups relative to protected, vaccinated groups (BCG+Adj, BAA). It should be emphasized that detection of these cytokines from pulmonary CD4^+^ T cells did not involve ex vivo stimulation of the cells. Thus, the cytokine expression levels observed likely closely mirror expression levels occurring in vivo.

Overall, here we report some key differences in the pulmonary immune cell populations induced by a highly protective TB vaccine using both latent and acute infection mouse models compared to less protective vaccines and nonprotected mice. The findings highlighted in this study may help lead to the design of a more protective vaccine for investigation in future nonclinical and clinical studies.

## ACKNOWLEDGEMENTS

This work was supported by FDA intramural funding. This research received no specific grant from any funding agency in the public, commercial, or not-for-profit sectors.

The authors declare no conflicts of interest.

The following reagent was obtained through BEI Resources, NIAID, NIH: ESAT-6, Recombinant Protein Reference Standard, NR-49424.

## Supplemental Material

**FIG S1.**
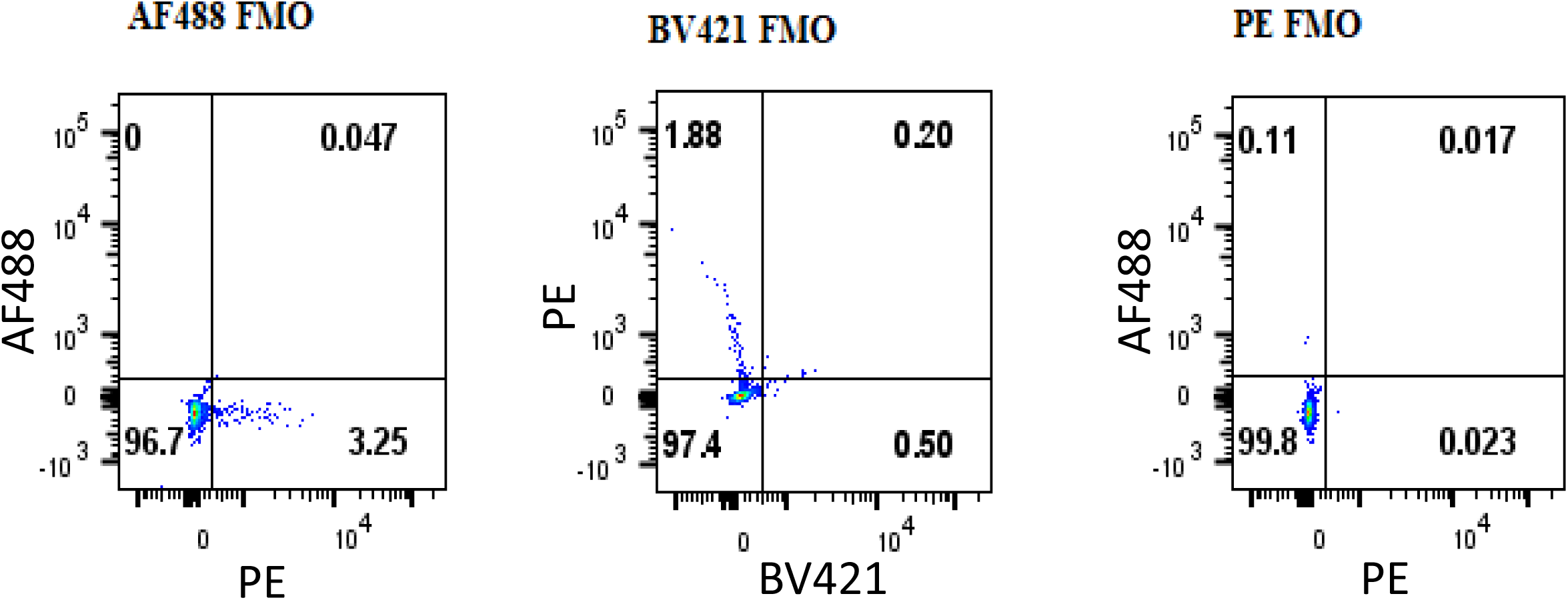
Representative dot plots showing FMO controls for gating on positive cell populations. Fluorescence minus one (FMO) control dot plots were used for placement of gates for determining frequencies of the different cell populations. These controls were created at the same time lungs were collected from infected mice for CFU determination and lung cell isolation. Antibody, fluorochrome combinations were as follows: TCRβ (AF700; clone H57-597), CD4 (PerCP-Cy5.5; clone RM4-5), IFN-γ (AF488; clone XMG1.2) and IL-17A (PE; clone TC11-18H10) and TGFβ (BV421; clone TW7-16B4).

**FIG S2.**
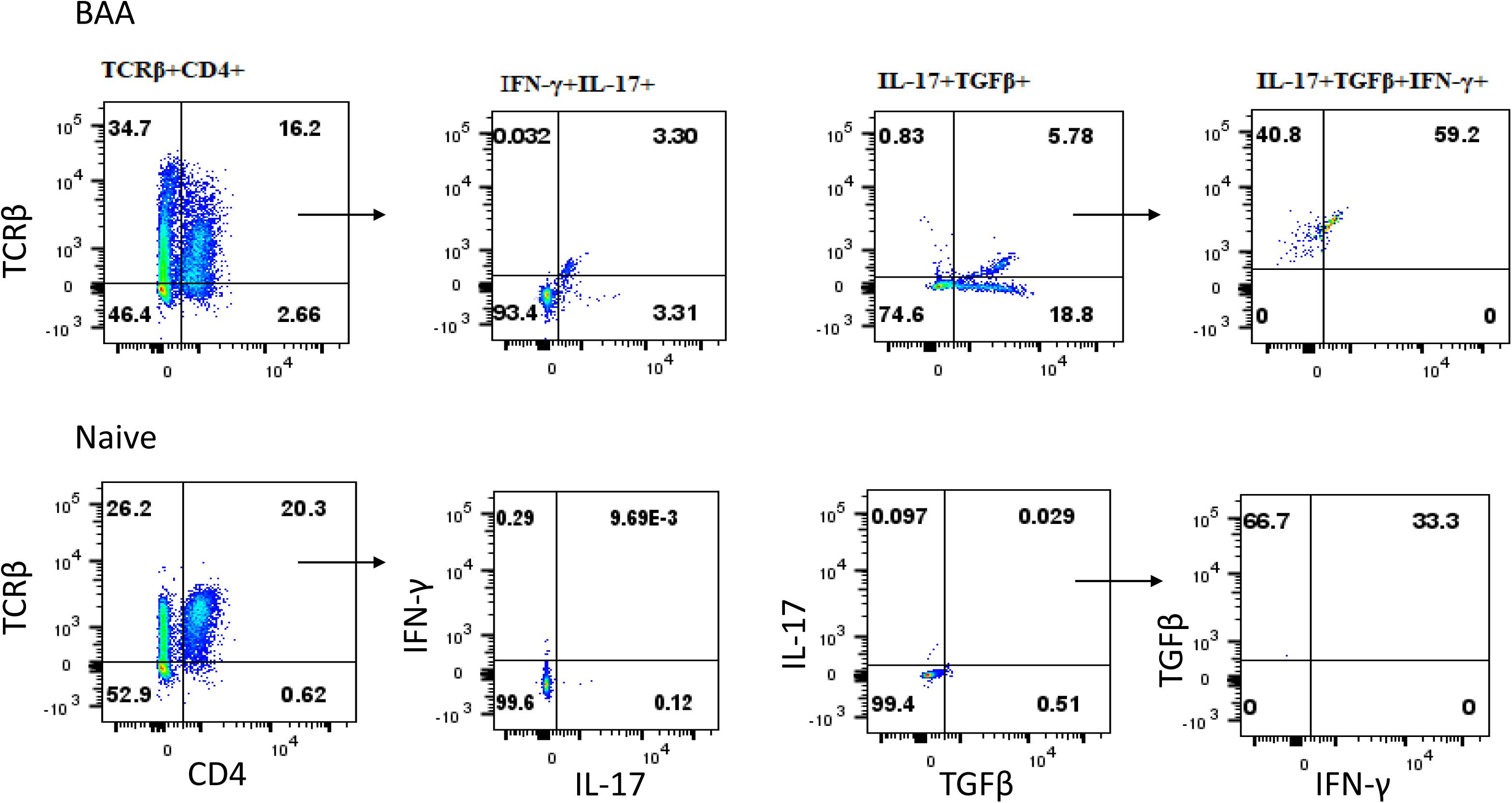
Representative dot plots showing the gating strategy for assessing the frequencies of monofunctional and multifunctional CD4+ T cells. A representative dot plot flow cytometry gating scheme is shown for determining frequencies of CD4^+^ T cells expressing only IFN-γ, TGFβ or IL-17A, or both IFN-γ and IL-17A or cells expressing all three cytokines from the lungs one month following an aerosol *M. tuberculosis* challenge with around 100 CFU. Antibody, fluorochrome combinations were as follows: TCRβ (AF700; clone H57-597), CD4 (PerCP-Cy5.5; clone RM4-5), IFN-γ (AF488; clone XMG1.2), IL-17A (PE; clone TC11-18H10) and TGFβ (BV421; clone TW7-16B4).

